# A Genetically Encoded System to Quantify and Evolve RuBisCO-Catalyzed Carbon Fixation

**DOI:** 10.1101/2024.05.15.594374

**Authors:** David L. Lanster, Zhiyi Li, Ahmed H. Badran

## Abstract

Strategies to study and alter the biochemical properties of RuBisCO often couple CO_2_ fixation to bacterial growth. However, these viability-coupled strategies are not quantitative and are limited by toxicity of the RuBisCO substrate RuBP, the slow kinetics of RuBisCO, and differences in RuBisCO expression. We report the development of the first genetically encoded system capable of accurately quantifying RuBisCO-dependent CO_2_ fixation *in cellulo* using a tripartite approach that combines bacterial strains which insulate RuBisCO-derived products, an engineered pathway that eliminates RuBP toxicity, and biosensors to concurrently monitor RuBisCO abundance and catalysis. We extended this biosensing strategy to 43 Form II and II/III RuBisCO homologs, finding strong agreement with *in vitro*-derived enzyme kinetics in living cells for the first time. Finally, we show how this system can be used to rapidly evolve functional RuBisCO enzymes. Our approach overcomes prior limitations by streamlining intracellular RuBisCO analyses and will enable the development of enzymes with improved CO_2_ fixation capabilities.

## Introduction

Carbon flux through the biosphere is mediated predominantly by the enzyme ribulose-1,5-bisphosphate carboxylase/oxygenase (RuBisCO), which catalyzes the capture and fixation of CO_2_ in the first committed step of the Calvin-Benson-Bassham (CBB) cycle. With global interest in sustainable strategies to reverse climate change, there has been increased focus on improving the CO_2_ fixation characteristics of RuBisCO. Indeed, as one of the most abundant proteins on Earth^1^, RuBisCO-dependent fixation accounts for >90% of all globally captured CO_2_, totaling ∼120 Gt CO_2_ per year^2^. However, the carboxylation rate of RuBisCO hinders photosynthetic efficiency: >95% of characterized RuBisCO homologs capture only 1–10 CO_2_ molecules per second^3^. Beyond slow carboxylation kinetics, RuBisCO can capture O_2_ instead of CO_2_, leading to the formation of the byproduct 2-phosphoglycolate (2PGL) that must be detoxified in photosynthetic organisms through the energy-intensive and CO_2_-emitting photorespiration pathway^4^. Photorespiration decreases net photosynthetic efficiency by up to 50%, returning a total of ∼59 Gt CO_2_ to the atmosphere annually^5,6^. While the promiscuous specificity of RuBisCO can be offset through CO_2_-concentrating mechanisms (CCMs) to minimize carbon loss, many photosynthetic organisms devote as much as 50% of their soluble protein towards biosynthesizing RuBisCO to support sufficient metabolic flux for biomass generation, making it one of the most abundant proteins on Earth^7^.

These poor biochemical characteristics of RuBisCO have made it a leading candidate for optimization to increase photosynthetic efficiency, which would have long-term implications for agricultural crop yields and as a platform for carbon capture biotechnologies^6^. Unfortunately, only modest gains have been realized through mining natural sequence diversity^8^, ancestral protein reconstruction^9–11^, and directed evolution^12–15^. State-of-the-art intracellular strategies to improve CO_2_ fixation rely on RuBisCO-dependent *Escherichia coli* (RDE) strains^12–16^, where RuBisCO-produced 3-phosphoglycerate (3PG) complements deletion of an essential glycolytic enzyme (e.g., GapA, Pgk). However, RDE strains have impaired growth rates^14^, a high false-positive rate^17^, and can require extensive *in vitro* characterization to confirm RuBisCO improvement^17,18^. In addition, many RDE-evolved RuBisCO alleles have been shown to improve solubility, folding, and/or expression rather than CO_2_ capture and fixation efficiency^12,15^. Driven by these limitations, robust methods are needed to quantify RuBisCO activity and abundance in living cells while concomitantly minimizing cell viability defects and selection escape.

We report the development of the first genetically encoded system capable of accurately quantifying RuBisCO-dependent carbon fixation *in cellulo*. We overcome the low throughput and slow growth rate of RDE approaches by decoupling RuBisCO activity from metabolism, which we achieve by engineering *E. coli* to use an underexploited glycolytic bypass that eliminates the accumulation of 3PG from glycolysis. Using this metabolically insulated chassis bacterium, we create an artificial biosynthetic pathway that controls the abundance of the toxic RuBisCO substrate ribulose 1,5-bisphosphate (RuBP) and converts RuBisCO-derived 3PG to glycerate, which we quantitatively monitor using an evolved transcriptional biosensor. Since RDE approaches can incentivize the evolution of increasingly soluble RuBisCOs, we developed a strategy to fine-tune and quantify intracellular RuBisCO abundance, which can be used to normalize biosensor activities. We extend these resources to 43 diverse RuBisCO enzymes to find good agreement between *in cellulo* biosensor activities and *in vitro*-derived kinetic parameters. Finally, we demonstrate the rapid enrichment of a functional RuBisCO enzyme from a library of inactive variants in only a few days. By showcasing how RuBisCO activity can be readily monitored and evolved in a cellular setting, we establish a high-throughput approach to quantify CO_2_ fixation and provide an effective platform for the directed evolution of RuBisCO enzymes with enhanced CO_2_ capture efficiencies.

### Metabolic Rewiring Insulates RuBisCO Products

RuBisCO captures CO_2_ by catalyzing the formation of a new carbon-carbon bond with RuBP, where the product spontaneously breaks down to yield two molecules of 3-phosphoglycerate (3PG). Since 3PG is a central metabolite in the *E. coli* Embden-Meyerhof-Parnas (EMP) pathway (**Extended Data Fig. 1a**), RDE approaches use this relationship to evolve RuBisCO: heterologously expressed phosphoribulokinase (Prk) generates RuBP, which RuBisCO carboxylates and cleaves to generate two molecules of 3PG in an EMP-deficient *E. coli* strain (**Extended Data Fig. 1b**)^13^. Rather than relying on this complementation strategy, we sought to insulate RuBisCO-derived products from host-encoded metabolism (**Extended Data Fig. 1c**) to enable the development of dedicated and quantitative biosensors for CO_2_ fixation.

We first consulted the STRING^19^ and EcoCyc^20^ databases to nominate proteins known or predicted to produce, consume, transport, and/or bind metabolites with structural or chemical similarity to 3PG or the product of oxygen fixation, 2-phosphoglycolate (2PG). This analysis yielded 39 candidate essential and non-essential genes for deletion^21^ (**Extended Data Table 1**). We first pursued deletions of all high priority non-essential genes in *E. coli* strain S1021^22^ using a λ-red recombineering strategy that permits positive–negative selection for scarless genome editing^23^ (**Extended Data Table 2**, **Extended Data Fig. 2**). This strategy gave rise to *E. coli* strain S3710, which grew comparably to S1021 (**Extended Data Table 3**). Next, we explored deletions of the relevant (and essential) EMP genes: *gapA*, *pgk*, *gpmA*, and *gpmM*. *E. coli* mutants lacking these genes can have reduced fitness and are often sensitive to glucose due to glycolytic metabolite buildup^24^, but remain viable using minimal media supplemented with glycerol and succinate (e.g., M9GS) ^25^. Using M9GS and S3710 cells, we deleted *pgk* to afford the *E. coli* strain S3711 (**Fig. 1a**) and confirmed its sensitivity to glucose (**Extended Data Fig. 3a**).

**Fig. 1.**
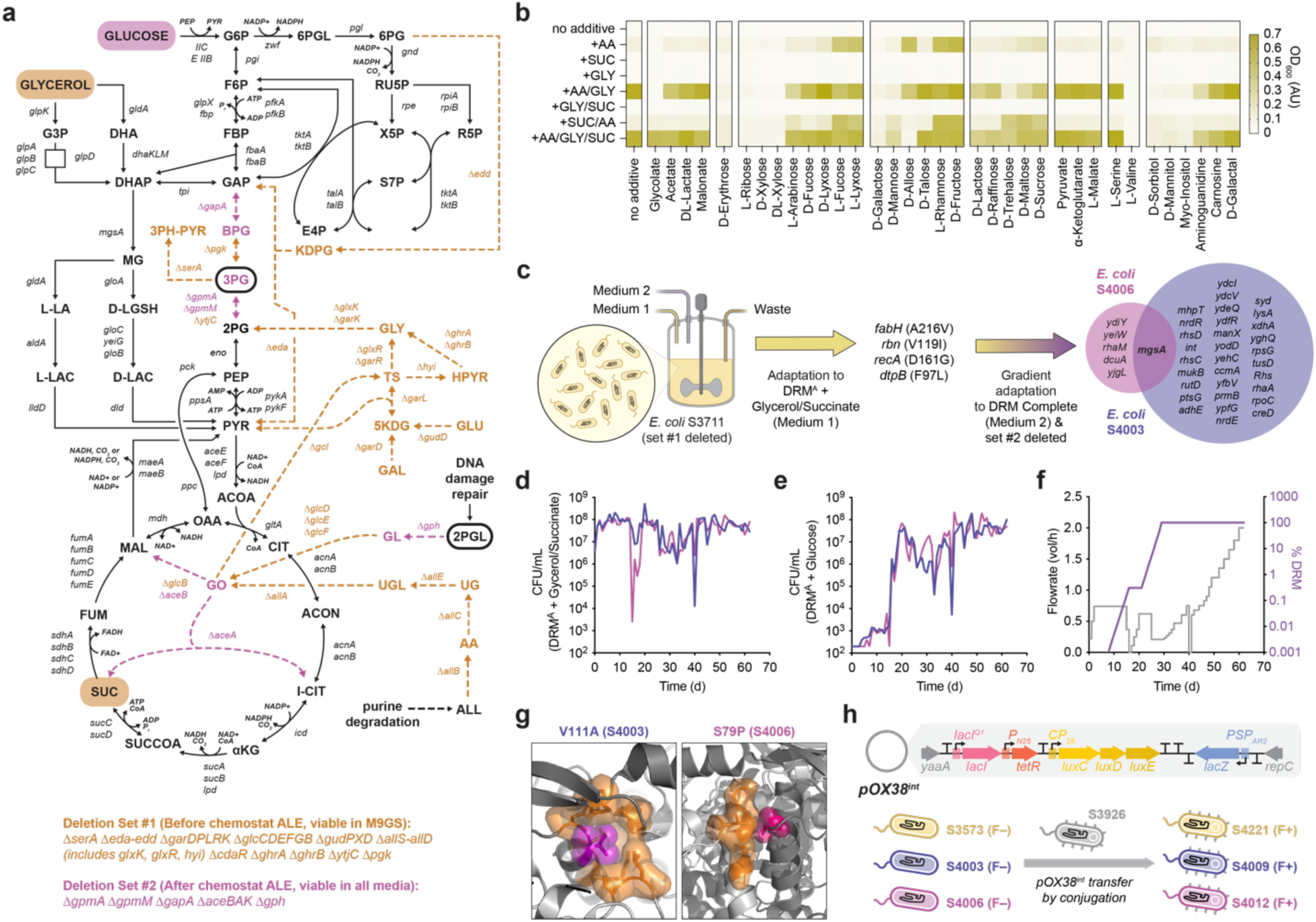
Metabolic Engineering and Adaptive Laboratory Evolution of RuBisCO-Insulated *E. coli* Strains. **a**, Engineered metabolism to insulate RuBisCO-derived 3PG and 2PGL (circled). **b**, Analysis of M9GS media supplements (10 mM) to improve S3711 fitness. Additives are grouped as follows: carboxylates, tetroses, pentoses, hexoses, di- and tri-saccharides, TCA intermediates, amino acids, and other metabolites. **c**, Adaptive laboratory evolution (ALE) schedule and whole-genome mutational analysis of S3711 cells over 62 days: neutral drift using DRM^A^ with glycerol and succinate (7 days), followed by complete DRM titration (22 days), then increasing chemostat flowrate (33 days). Final ALE strains S4003 and S4006 share mutations in MgsA only. **d**, Cell titer throughout the ALE campaign quantified on DRM^A^ with glycerol and succinate. **e**, Cell titer throughout the ALE campaign quantified on complete DRM. **f**, Schedule of chemostat flowrate and medium composition. Flowrate reductions indicate short pauses to recover cell densit. **g**, MgsA crystal structure (PDB: 1B93; ^61^) highlighting the homohexamer interfacial mutations V111A (strain S4003) and S79P (strain S4006). Mutations are shown in purple/violet, and packing interface in orange. **h**, A pOX38^Tc^-derived ^31^ mobilizable F’ plasmid called pOX38^int^, which integrates genes for LacI, TetR, LuxCDE, and LacZ between the *yaaA* and *repC* genes. The engineered strain S3573 (S3711 with native *pgk*) and evolved strains S4003 and S4006 were conjugated with the pOX38^int^ donor S3926 to yield the S4221, S4009, and S4012 strains, respectively. All datasets include 1-2 biological replicates.

Further genome editing of S3711 cells was not possible, perhaps reflecting unknown metabolic dependencies in M9GS. However, no alternative medium formulation improved S3711 fitness beyond M9GS (**Fig. 1b**). We therefore evolved S3711 cells through adaptive laboratory evolution (ALE) to restore its fitness and enable additional genetic deletions, and chose to use Davis Rich Medium (DRM)^22^ since it has been previously used to effectively grow *E. coli* in chemostat settings with optimal resource economy (**Fig. 1c**). We first cultured S3711 cells in DRM^A^ (phosphate-buffered amino acids and nucleobases) with glycerol and succinate to incentivize genetic drift, which we envisioned would enable the discovery of mutations with improved fitness in glucose-rich medium downstream (**Fig. 1c**). In order to minimize the stringency of this selection, we titrated complete DRM (contains 25 mM glucose)^22^ at an initial concentration of 0.001% to maintain the glucose concentration below the 1 µM toxicity threshold^26^. We progressively doubled the DRM concentration to 100% over 22 days, and then increased the flow rate to 2 vol h^-1^ over 33 days to ensure that evolved cells could readily grow in this medium at an appreciable rate (**Fig. 1d**). Following daily sampling throughout the 62-day ALE campaign, we quantified cell titer on DRM^A^ with glycerol/succinate (**Fig. 1e**), DRM (**Fig. 1f**), and the standard laboratory medium 2xYT (**Extended Data Fig. 3b, c**). Interestingly, both chemostat populations showed a marked fitness increase in DRM and 2xYT at 13-14 days (**Extended Data Fig. 3d, e**).

Chemostat populations at 62 days were used for subsequent genome modification to preserve allelic diversity. We deleted all remaining candidate glycolytic enzymes as well as *gph* and *aceBAK* (to eliminate glycolate and glyoxylate production, respectively) from the *E. coli* genome, corresponding to 31 genes across 46,640 base pairs in total. We noted that these deletion efforts culminated in a phenotypically distinct strain from each chemostat population: S4003 and S4006. Whole genome sequencing revealed that each isolate independently converged on a unique mutation to methylglyoxal synthase (MgsA), the rate limiting enzyme in the methylglyoxal bypass (**Extended Data Table 4**, **Fig. 1c**). Whereas methylglyoxal is a toxic electrophile that can irreversibly modify proteins^27^, it is thought to affect the transition the cell’s metabolic profile between conditions of resource abundance and starvation^28^. Both mutations in *mgsA* map to the homohexamer interface (**Fig. 1g**), which regulates allostery and cooperativity during catalysis^29^. Since both strains can robustly grow in medium containing glucose but did not acquire mutations predicted to limit glucose uptake (**Extended Data Table 4**), we speculate that the methylglyoxal pathway now supports an effective glycolytic bypass, a hypothesis corroborated by a recent finding that MgsA overexpression permitted growth on glycerol in an EMP-deficient *E. coli*^30^. Finally, we completed our strain development efforts by engineering an F’ plasmid, pOX38^int^ (based on pOX38::Tc)^31^, that expresses key proteins that would streamline our reporter assays: the transcriptional repressors LacI and TetR for tunable promoter control, LuxCDE to generate the LuxAB substrate decanal *in situ*, and the membrane integrity-responsive LacZ cassette^32^ (**Fig. 1h**). We mated the key strains S3711, S4003, and S4006 with the pOX38^int^ donor strain S3926 to create S4221, S4009, and S4012, respectively (**Extended Data Table 5**). These mated strains were used for all subsequent assays.

### Glycerate-Responsive Biosensor Development

Whereas RDE systems couple CO_2_ fixation to cell fitness^12–15^, we aimed to monitor RuBisCO product formation using a biosensor strategy to streamline future characterization of CO_2_ capture efficiency. While no known biosensors respond to 3PG, the transcriptional activator CdaR responds to its dephosphorylated counterpart glycerate^33,34^ (**Fig. 2a**). We reasoned that it may be combined with a phosphatase to quantify 3PG production by RuBisCO. We tested CdaR-responsive promoters^33,34^ in S4221 cells alongside CdaR overexpression from a second plasmid, finding that P_garP_ responded most sensitively to glycerate (**Fig. 2b**). While this dynamic range could be improved by condensing the genetic circuit to a single plasmid in S4221 cells (**Fig. 2c, d**), P_garP_ was constitutively activated in the evolved S4009 and S4012 strains following ALE metabolic optimization (**Fig. 2d**). We hypothesized that CdaR was activated by an unknown off-target metabolite following ALE, and thus developed a dual positive-negative selection scheme to evolve CdaR variants with greater specificity for glycerate (**Fig. 2e**). After one round of error-prone PCR and selection, we uncovered CdaR mutants with glycerate-dependent reporter fluorescence (**Fig. 2f**), where isolates evolved in DRM outperformed those from 2xYT (**Fig. 2g**). Sequencing of the best performing clones from DRM and 2xYT revealed convergence on mutations predicted to affect a putative interfacial ligand-binding pocket (**Fig. 2h**). The DRM-derived consensus variant CdaR^evo1^ reestablished a large glycerate-dependent dynamic range in S4012 cells across multiple reporter contexts (**Fig. 2i, j**)^35–37^, nominating this mScarlet-I biosensor–strain combination as the leading chassis for further biosensor development.

**Fig. 2.**
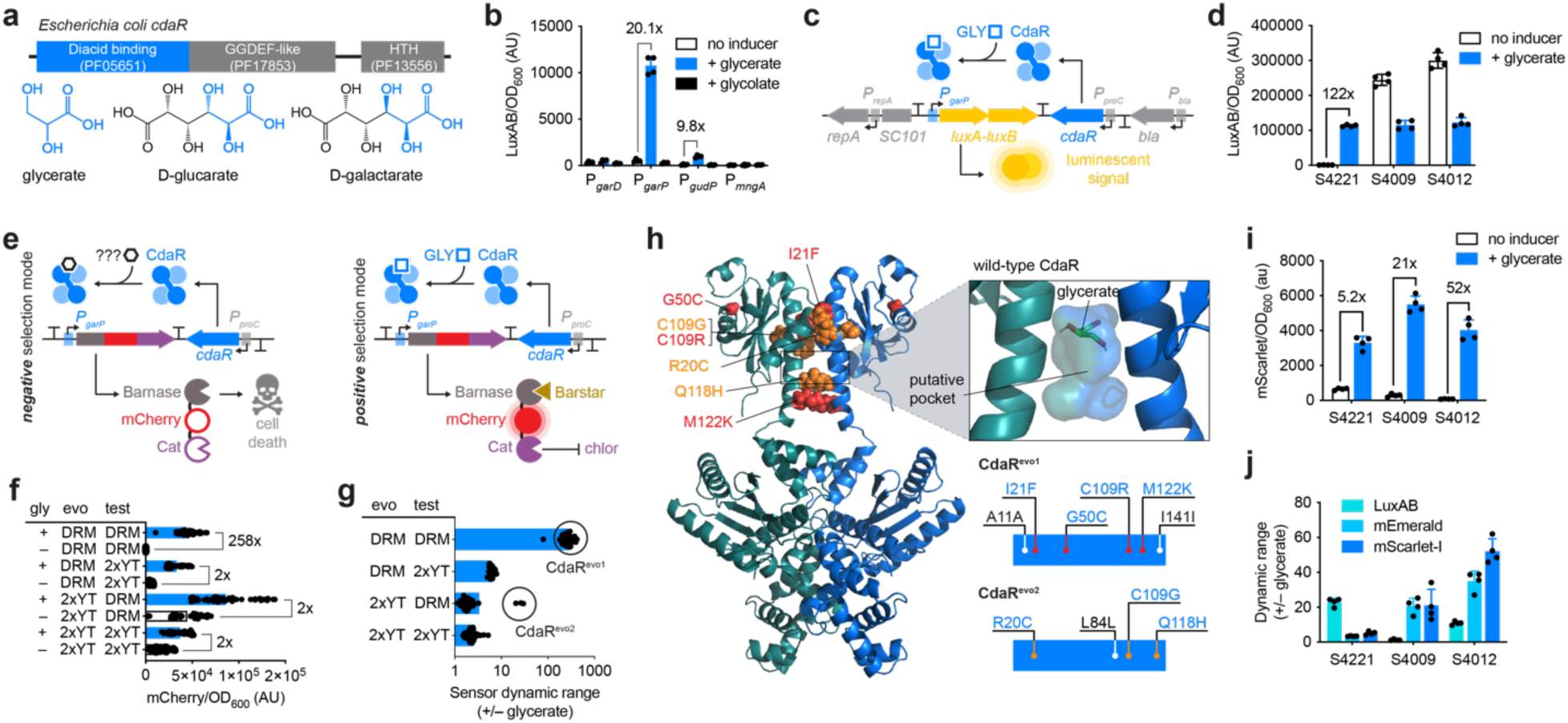
Development of a Glycerate-Specific Biosensor Compatible with Metabolically Insulated Strains. **a**, Domain organization of the carbohydrate diacid regulator CdaR. Diacid binding domain and shared chemotypes between glycerate, D-glucarate, and D-galactarate are shown in blue. **b**, Testing CdaR-regulated promoters using glycerate and glycolate, nominating the glycerate-specific P_garP_ with the greatest signal and dynamic range. **c**, Schematic representation of the optimized glycerate-responsive biosensor: CdaR is constitutively expressed and requires glycerate (GLY) to activate P_garP_ transcription, resulting in a luminescent signal through LuxAB. Grayscale elements correspond to the origin of replication and resistance marker. **d**, The single plasmid glycerate-responsive biosensor improves dynamic range in S4221 cells, but is constitutively activated in the evolved strains S4009 and S4012, suggesting unintended crosstalk with an ALE-derived cellular metabolite. **e**, Selection strategy to improve CdaR substrate specificity in S4012 cells: negative selection (without glycerate): undesired promoter activation produces the toxin barnase and results in cell death. Positive selection (with glycerate): barnase is inhibited by the constitutively expressed antitoxin barstar, and glycerate-dependent activation produces a fluorescent signal (mCherry) and chloramphenicol resistance (Cat). **f**, Selection outcomes depend on the medium used for evolution (evo) and subsequent assays (test). Directed evolution of CdaR in DRM yielded variants with greatest biosensor activation. The labels “evo” and “test” indicate the medium used for directed evolution and the medium used in the shown assay, respectively. **g**, Single clone analysis and Sanger sequencing of variants with large dynamic range yielded two genotypes: Cdar^evo1^ and Cdar^evo2^. **h**, Mutational mapping of evolved mutations onto an Alphafold-predicted ^62,63^ CdaR dimer shows that mutations may affect a putative ligand binding pocket at the dimer interface. Glycerate docked using CB-Dock ^64,65^. Evolved mutations are shown schematically in the CdaR diacid binding domain. Red (Cdar^evo1^) and orange (Cdar^evo2^) mutations are coding, whereas mutations in white are non-coding. **i**, Cdar^evo1^ restores glycerate-dependent biosensor activity in the metabolically insulated strains S4009 and S4012. **j**, Cdar^evo1^ benchmarking using various strains and reporters. S4012 cells encoding Cdar^evo1^ driving an mScarlet-I reporter were used for all subsequent assays. All datasets include 4 biological replicates.

### Broad-Scope Phosphatases Convert 3PG to Glycerate

CdaR^evo1^-dependent detection of RuBisCO activity requires dephosphorylation of 3PG to yield glycerate, yet no phosphatases that act on 3PG *in vivo* have been reported to date. Since exogenous 3PG cannot traverse cell membranes^38^, we expressed *S. typhimurium* PgtP (*St*PgtP) ^38^ and *E. coli* UhpT_D388C_ (*Ec*UhpT_D388C_)^39^ to import exogenous 3PG into cells (**Fig. 3a**). To establish our approach we tested inducible phosphatases that are known to dephosphorylate 3PG *in vitro: E. coli* Gph, YieH, and YbhA^40^; *Saccharomyces cerevisiae Sc*Pho13^41^ and *Sc*YOR283w ^42^; *Thermus thermophilus Tt*HA0368^43^; *Geobacillus stearothermophilus Gs*PhoE^43^. Whereas Gph and *Gs*PhoE non-specifically activated CdaR^evo1^ (**Fig. 3b**), the remaining enzymes Gph, YbhA, *Sc*YOR283w, *Tt*HA0368, and *Gs*PhoE all activated CdaR^evo1^ in a 3PG-dependent manner (**Fig. 3c, d**), with *Ec*UhpT_D388C_ outperforming *St*PgtP. Constitutive expression of these five phosphatases prior to 3PG incubation further amplified CdaR^evo1^-dependent dynamic range (**Fig. 3e, f**). To nominate the most effective phosphatase, we titrated the top performing enzymes YbhA, *Sc*TOR283w, and *Tt*HA0368 with glycerate and 3PG. YbhA and *Sc*TOR283w activated CdaR^evo1^ to a greater extent than the no-phosphatase control across all glycerate concentrations, suggesting an unintended false-positive impact on biosensor performance. This effect was consistent in the 3PG titration, whereas *Tt*HA0368 showed the predicted response in both settings (**Fig. 3g, h**). We confirmed the on-target activity of *Tt*HA0368 by nominating putative catalytic residues through homology and structural modeling (**Extended Data Fig. 4a-c**) and show that mutating active site residues abolishes phosphatase activity and CdaR^evo1^-dependent signal generation (**Fig. 3i**). These results establish that CdaR^evo1^ and *Tt*HA0368 can be used together to quantify intracellular 3PG concentrations in metabolically-insulated *E. coli* strains.

**Fig. 3.**
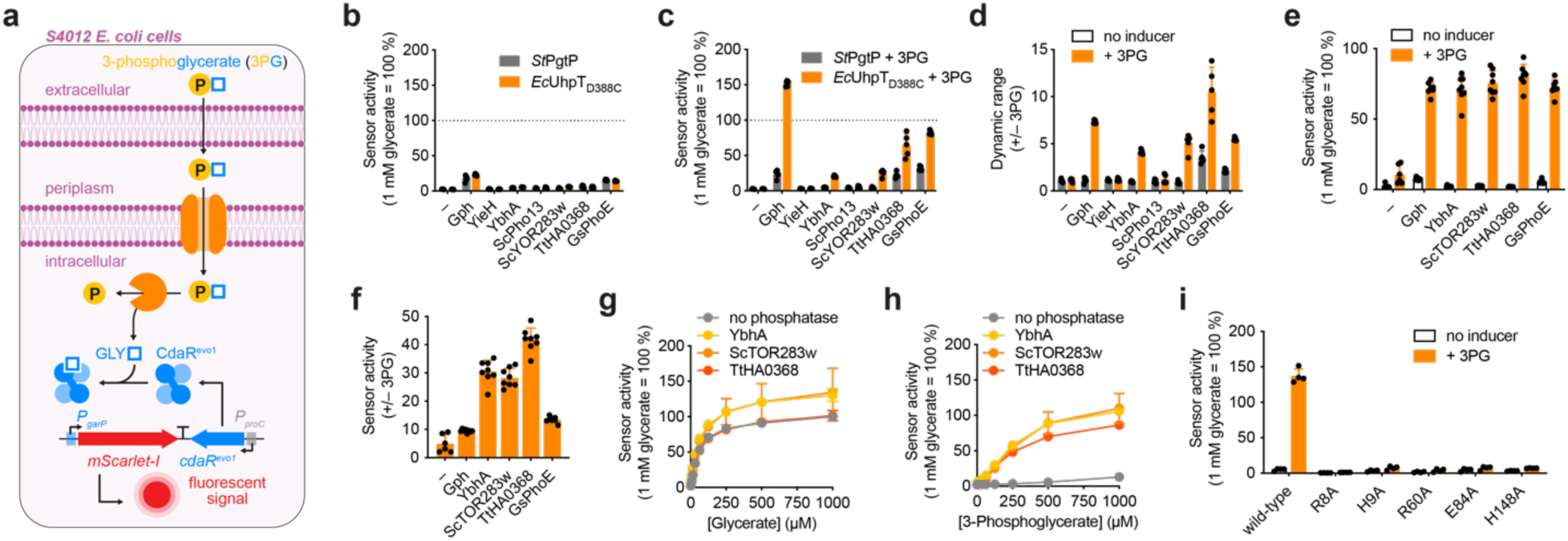
Discovery of a 3PG-Permissive Phosphatase to Generate Glycerate in Living Cells. **a**, Strategy for phosphatase and importer analysis. Exogenous 3PG is transported into the cytosol by an inner membrane transporter. The phosphatase cleaves 3PG to yield glycerate, which is detected by CdaR^evo1^. **b**, Spurious activation of the CdaR^evo1^ biosensor by Gph and *Gs*PhoE in the absence of exogenous 3PG. **c**, On-target CdaR^evo1^ biosensor activation after exogenous 3PG addition in a phosphatase-dependent manner. *Ec*UhpT_D388C_ outperformed *St*PgtP alongside all tested phosphatases in response to 3PG. **d**, CdaR^evo1^ biosensor dynamic range +/– 3PG using all phosphatases and importers in response to 3PG. **e**, Low-level constitutive phosphatase expression improves biosensor signal in response to 3PG. **f**, Low-level constitutive phosphatase expression improves biosensor dynamic range in response to 3PG, spearheaded by YbhA, ScTOR283w, and TtHA0368. **g**, Glycerate titration analysis in S4012 cells encoding the top performing phosphatases, highlighting the undesirably high CdaR^evo1^ biosensor activation when using YbhA and *Sc*TOR283w. **h**, 3PG titration analysis in S4012 cells, showcasing phosphatase- and 3PG-dependent CdaR^evo1^ biosensor activation. Dose-dependent signal increase without phosphatase likely reflects spontaneous 3PG dephosphorylation prior to cell entry or dephosphorylation by periplasmic phosphatase(s). **i**, Inactivating mutations to the putative catalytic residues of *Tt*HA0368 ablate CdaR^evo1^ biosensor activation. All datasets include 4-8 biological replicates.

### Kinetic Tuning Alleviates Prk Toxicity

The RuBisCO substrate RuBP is not natively biosynthesized by *E. coli,* which requires the expression of phosphoribulokinase (Prk) and RuBisCO to complete the Calvin Benson Bassham cycle. Heterologously expressed Prk can phosphorylate ribulose-5-phosphate (R5P), a pentose phosphate pathway intermediate, to generate RuBP in *E. coli*^13^. However, Prk activity is toxic to *E. coli* and results in frequent selection escape in RDE strains (**Fig. 4a**)^12–15^. We hypothesized that this toxicity is exacerbated by efficient, high-level expression Prk constructs used in prior reports (**Fig. 4b**), which may be further confounded by stochastic induction of the P_BAD_ promoter in cells lacking constitutive arabinose importer expression^44^. We addressed these issues by reducing *Synechococcus elongatus* Prk expression in *E. coli* using the low-copy RK2 origin^45^, the tightly controlled rhamnose-dependent promoter P_rha_^46^, and a weakened RBS series^47^ (**Fig. 4b**), but found that all designs still induced toxicity in S4012 cells (**Fig. 4c**). We hypothesized that mutations to the ATP or R5P binding sites (**Fig. 4d**)^48^ could kinetically tune Prk activity to eliminate toxicity while still providing sufficient RuBP for RuBisCO. To concurrently investigate both points, we combined the *Rhodospirillum rubrum* RuBisCO (CbbM), *R. rubrum* carbonic anhydrase (CA)^49^, *Tt*HA0368, and CdaR^evo1^ biosensor with Prk mutants^48^ driven by a weak promoter, then monitored biosensor signal generation under ambient (0.04%) and high CO_2_ (5%) conditions to confirm full pathway activity (**Fig. 4e**). The CO_2_ concentration-dependent biosensor signal from this 5-plasmid genetic circuit confirmed full pathway activity and showed that Prk mutants R52A and W140A were well tolerated (**Fig. 4f**).

**Fig. 4.**
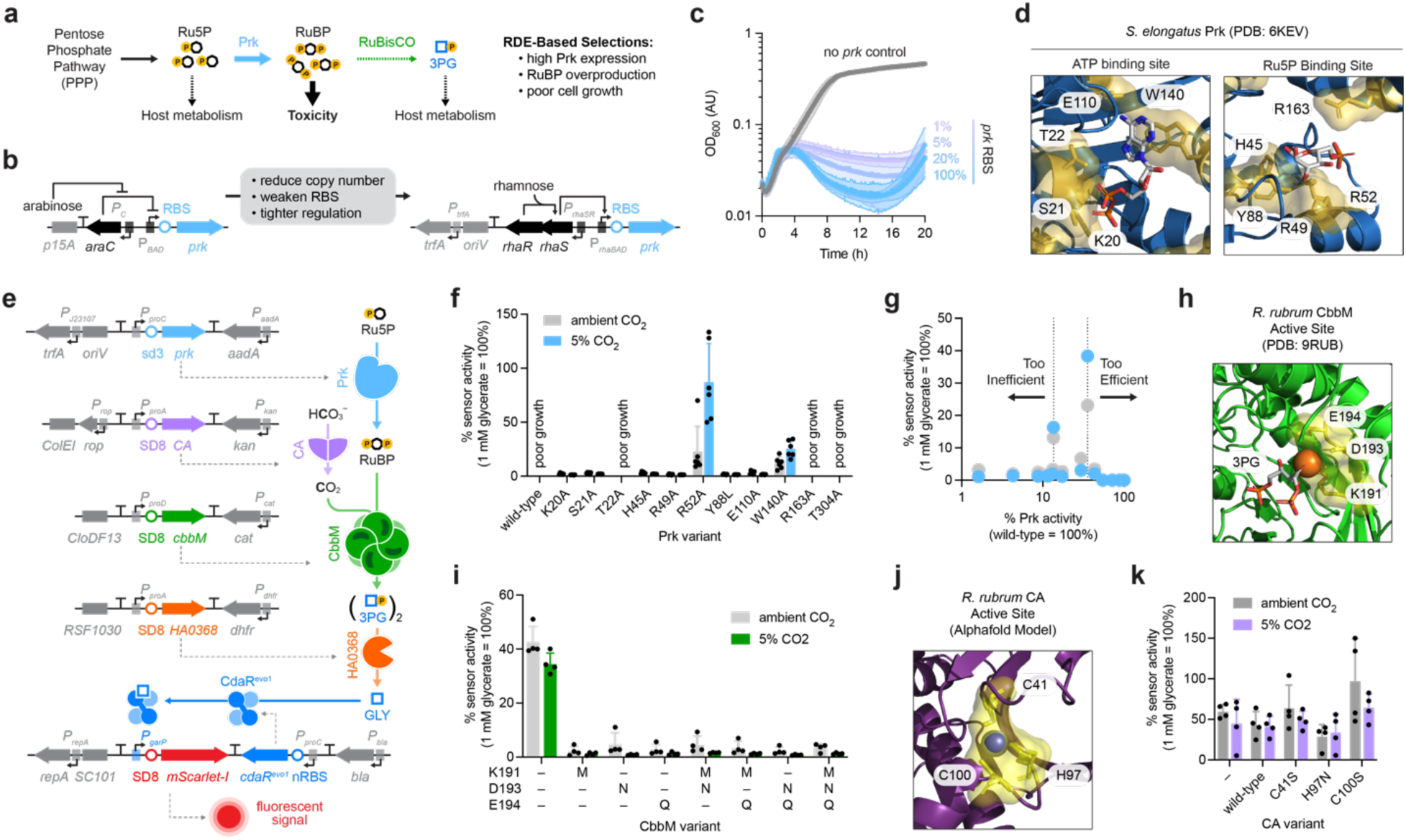
Genetic and Biochemical Prk Tuning Eliminates Toxicity and Enables Biosensor-Dependent RuBisCO Monitoring in Living Cells. **a**, Concentration-dependent RuBP toxicity and false-positive outcomes in RDE strains may result from unnecessary Prk overexpression and insufficient detoxification by RuBisCO. **b**, A new Prk expression plasmid that encodes the low copy number RK2 origin ^45^, weakened RBSs ^66^, and tighter transcriptional control through the rhamnose promoter ^67^. **c**, Kinetic analysis of S4221 cell density, showing Prk concentration-dependent toxicity despite plasmid engineering. Cells were induced with 0.1 mM L-rhamnose at t = 0 hours. **d**, Target residues for Prk mutagenesis (yellow) in the ATP and Ru5P binding sites. **e**, Schematic representation of the complete RuBisCO biosensor in S4012 cells: Prk (or active site mutant thereof) converts Ru5P to RuBP, which is then combined with carbonic anhydrase (CA)-derived CO_2_ by a functional RuBisCO to yield two molecules of 3PG. *Tt*HA0368 dephosphorylates 3PG to glycerate, resulting in CdaR^evo1^ biosensor activation. **f**, Comparison of the full RuBisCO biosensor pathway outcome using Prk active site mutants, showcasing a moderate CO_2_ concentration-dependent CdaR^evo1^ biosensor activation using Prk mutants R52A (Ru5P binding site) and W140A (ATP binding site). **g**, Comparison of CdaR^evo1^ biosensor activation to predicted Prk activity highlights a narrow window (15-45% wild-type activity) that affords a robust signal without adversely affecting S4012 cell viability. **h**, Structure of *R. rubrum* CbbM with essential catalytic residues used to inactivate RuBisCO highlighted (yellow). **i**, Mutation of single or multiple CbbM active site residues to alanine ablates biosensor activity, confirming that RuBisCO-dependent 3PG generation is necessary to achieve robust biosensor activation. **j**, *R. rubrum* carbonic anhydrase (CA) Alphafold model with the catalytically necessary Zn^2+^ ion and essential residues in the active site (yellow). **k**, Mutation of CA active site residues to alanine do not impact biosensor activity, suggesting that CO_2_ is not limiting under assay conditions (5% CO_2_) to achieve robust biosensor activation. All datasets include 4-22 biological replicates.

These results further established a narrow window of 15-45% wild-type Prk activity as necessary to provide sufficient RuBP for RuBisCO-dependent CO_2_ fixation without affecting cell viability (**Fig. 4g**). To confirm that the observed signal was dependent on a functional RuBisCO, we inspected the CbbM structure and nominated conservative mutations to key residues in the CbbM active site (**Fig. 4h**), where all mutants and combinations thereof ablated activity (**Fig. 4i**). Surprisingly, CA activity (which is often included in RDE strains to increase intracellular CO2 concentrations)^49^ was not necessary as active site mutations predicted to eliminate its catalytic activity did not influence biosensor readout (**Fig. 4j, k**). CA was therefore removed from downstream assays. Taken together, these results establish the first biosensor capable of monitoring RuBisCO-dependent CO_2_ fixation in a living cell.

### Systematic Medium, Strain, and Circuit Optimization

Having established our RuBisCO-dependent biosensor, we next sought to optimize our system for future analyses. We explored the medium dependence of our biosensor–strain combination, in part due to the outcomes of our CdaR evolution campaign (**Fig. 2f, g**). We found that S4012 cells carrying all 5 plasmids grew to different densities across common microbial media (**Extended Data Fig. 5a**), with unique impacts on biosensor activity and dynamic range (**Extended Data Fig. 5b, c**). By comparing outgrowth conditions that maximized cell density with assay conditions that retained biosensor dynamic range (**Extended Data Fig. 5d**), we identified that DRM supplemented with peptone, tryptone, or yeast extract greatly improved cell density, but also inadvertently activated the biosensor at high concentrations (**Extended Data Fig. 5e, f**). Using an unbiased medium optimization approach, we further uncovered that the phosphoenolpyruvate (PEP) transition-state analog oxalate^50^ was a promising additive that improved S4012 cell growth (**Extended Data Fig. 5g, h**). These optimizations together gave rise to enhanced DRM (eDRM), a DRM-based medium that is supplemented with peptone, yeast extract, and oxalate at specific concentrations that maximize cell density without affecting biosensor accuracy or dynamic range (see Methods).

We next sought to validate *R. rubrum* CbbM activity on eDRM agar plates. Curiously, we noted spurious activation of the mScarlet-I signal in cells encoding the catalytically inactive CbbM mutant K191M when plated near cells expressing the functional wild-type allele. This finding suggested that glycerate may passively diffuse or be actively transported from a functional RuBisCO-encoding strain to a non-functional counterpart, which would result in a high false-positive discovery rate in future directed evolution studies. To establish the responsible transporter(s), we generated glycerate donor and acceptor strains that could be orthogonally monitored using mScarlet-I and mEmerald biosensors, respectively (**Extended Data Fig. 8a**). Next, we deleted transporters from the S4012 genome based on their reported ability to transport ligands with chemical similarity to glycerate (**Extended Data Table 6**) and tested them using an agar-based diffusion assay. This experiment confirmed that the lactate/glycolate permease LldP was responsible for glycerate entry in S4012 cells (**Extended Data Fig. 8b**). The resulting LldP-deleted strain S5643 (S4012 Δ*lldP::kan*) was used for all subsequent analyses. Finally, we condensed the *prk(R52A)* and *cdaR^evo1^* genes into a low-copy SC101 plasmid and cultured all strains henceforth in CO_2_-rich conditions without a CA expression plasmid, resulting in a three-plasmid biosensor system with reduced cellular burden.

### Quantitative RuBisCO Benchmarking and Enrichment

We extended this second-generation biosensor to 43 Form II and II/III RuBisCO homologs (**Fig. 5a**)^51^, which we hypothesized could be used to confirm *in vitro*-derived enzyme kinetics in living cells for the first time. Since each homolog may be differentially expressed *E. coli*^8,13^, we leveraged a split sfGFP tagging strategy to quantify RuBisCO concentration during analysis (**Fig. 5b**)^52^. Whereas most RuBisCO homologs activated the glycerate biosensor, mScarlet-I signal moderately correlated with *in vitro* activity (*r*=0.2795; *p*=0.0731) (**Fig. 5c**).

**Fig. 5.**
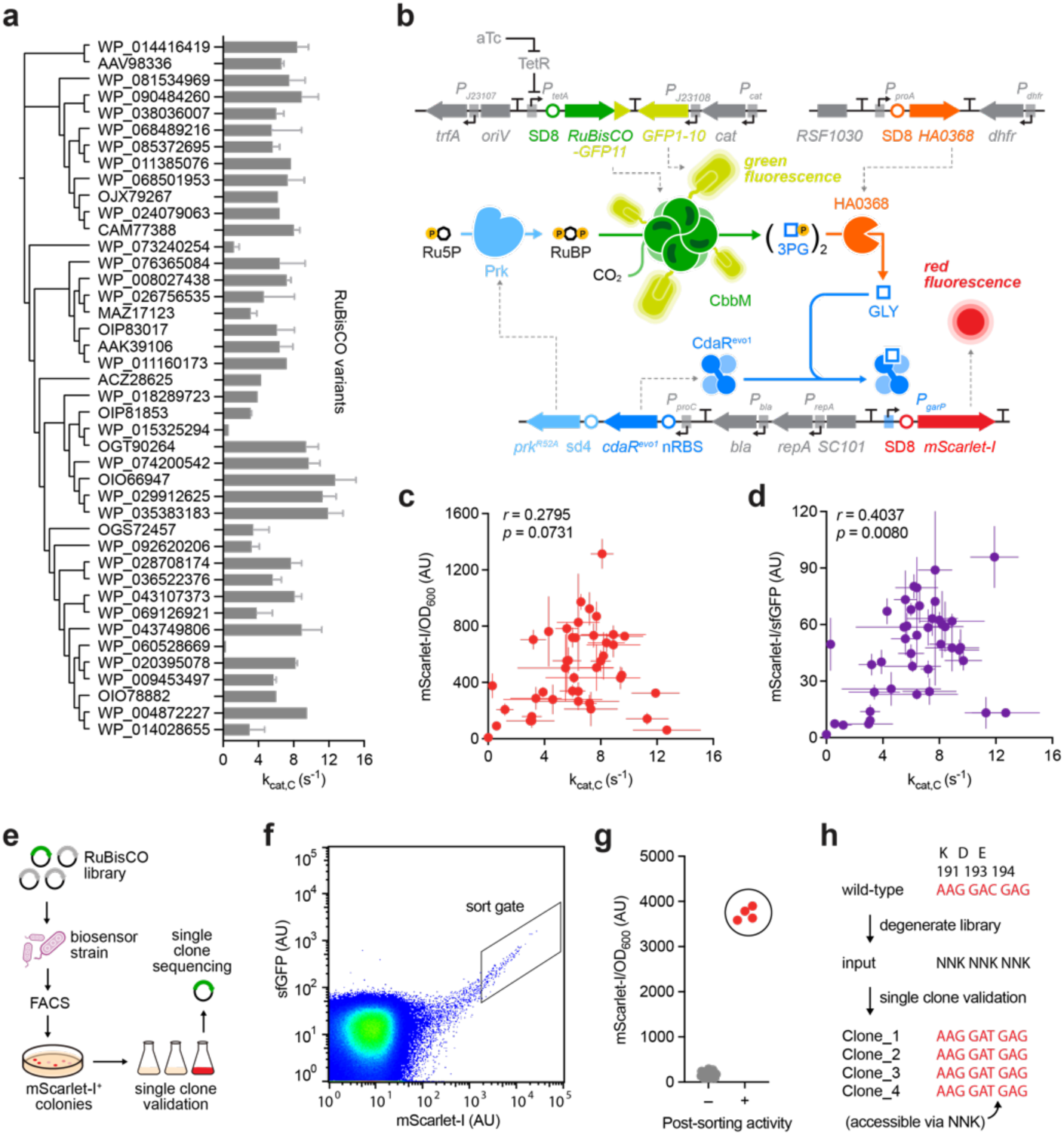
Intracellular Quantification of RuBisCO Activity and Enzyme Concentration Enables Rapid Validation of *In Vitro*-Derived Kinetic Parameters and Functional Enzyme Enrichment. **a,** Unrooted phylogenetic tree of 43 previously characterized Form II and II/III RuBisCO variants using an *in vitro* enzyme coupled assay ^8^. Bars indicate k_cat_ for CO_2_ capture and subsequent 3PG generation at 10% CO_2_. **b**, Strategy to quantify RuBisCO abundance in *E. coli*. Each RuBisCO variant includes a C-terminal sfGFP11 tag that binds to sfGFP1-10 to afford a fluorescent signal ^52^. RuBisCO-sfGFP11 expression is induced by aTc, whereas sfGFP1-10 is constitutively expressed. **c**, Comparison of CdaR^evo1^ biosensor activation using 43 RuBisCO variants to *in vitro* derived kinetics without normalization for RuBisCO abundance. **d**, After normalization to sfGFP signal (indicates RuBisCO abundance), agreement between CdaR^evo1^ biosensor activation and *in vitro* derived kinetics is improved to provide a more accurate quantification of intracellular bioactivity. **e**, Scheme for mock evolution of *R. rubrum* CbbM from a degenerate library of >32,000 variants. A library is generated by site-saturation mutagenesis of positions K191/D193/E194 and introduced into the biosensor strain. Following FACS, sorted colonies are plated on conventional agar plates and screened for an mScarlet-I^+^ phenotype, followed by single clone sequencing. **f**, FACS plot of sfGFP and mScarlet signals with gate denoting the sorted population. **g**, Single clone validation using the mScarlet-I biosensor, nominating the circled clones are positives. **h**, Sequencing of sorted colonies confirms the wild-type allele at all three positions. Whereas the original CbbM plasmid contains the aspartate codon GAC, all recovered clones contained the GAT aspartate codon which indicates their origin from the NNK library (GAC is not included in the library). All datasets include 3-4 biological replicates.

Normalization of biosensor readout by RuBisCO abundance (via sfGFP reassembly) resulted in an improved correlation with previously reported kinetic rate constants for CO_2_ capture (*r*=0.4037, *p*=0.0080) (**Fig. 5d**). Based on these findings, we envisioned that this biosensor may be used to evolve RuBisCO enzymes with altered properties. To demonstrate this, we used a site-saturation approach to create degenerate NNK libraries at the catalytic residues K191, D193, and E194 in the *R. rubrum* CbbM RuBisCO active site^53^ and attempted to enrich for the functional alleles via FACS (**Fig. 5e**). Prior biochemical analysis suggests that all three positions must converge on the wild-type amino acids to be functional, such that only a single variant in the >32,000-member library would be active. Following FACS (**Fig. 5f**) and confirmation of single clone activity via mScarlet-I fluorescence, we used single clone sequencing to confirm that all mScarlet-I positive cells had indeed re-discovered the correct identities at positions K191, D193, and E194 (**Fig. 5i**). These results correspond to >32,000-fold enrichment of a functional RuBisCO enzyme in a single round. Taken together, these results establish a quantitative, high-throughput strategy for RuBisCO directed evolution with a greatly diminished false-positive rate compared to traditional RDE approaches.

## Discussion

Improving CO_2_ fixation by RuBisCO has remained one of the most important directed evolution goals since its discovery nearly 70 years ago^54^. Recognizing the low-throughput and scarcity of genetic tools in plant-and algal-based directed evolution hosts^55,56^, Morell and colleagues proposed that complementation strategies could be used to experimentally investigate and evolve RuBisCO variants with improved catalytic properties in *E. coli*^57^. While RuBisCO-dependent CO_2_ fixation can support *E. coli* growth, applying selection pressure on organismal fitness aimed at improving a single gene can often have pleotropic consequences, including false positives^17^ and upstream pathway adaptation^49,58^. On the other hand, modern synthetic biology approaches to improve enzyme activity routinely make use of natural or engineered biosensors for directed evolution campaigns^59^. This framework has been challenging to apply to RuBisCO since its product, 3PG, is a central glycolytic metabolite necessary for *E. coli* viability. Rather than integrate RuBisCO into *E. coli* metabolism, we orthogonalized RuBisCO to allow for activity measurements and selection pressures in isolation from the host. After metabolic rewiring to eliminate 3PG production and catabolism, adaptive evolution to re-balance metabolic flux revealed an underappreciated glycolytic bypass^60^ (**Fig. 1a**), and permitted the creation of an artificial biosynthetic cascade and transcriptional biosensor for the quantitative monitoring of RuBisCO activity. Improving RuBisCO catalysis has been out of reach using traditional technologies, leading to the belief that RuBisCO may be evolutionarily trapped or that improvements are biophysically intractable. While the prevailing models suggest that improvements to RuBisCO will require extensive changes to this enzyme, the resources described herein (**Extended Data Tables 5, 7**) are poised to tackle these exciting evolution and bioengineering campaigns.

## Methods

### General methods

All PCRs were performed using Phusion U HotStart DNA Polymerase (Life Technologies), Q5U Hot Start High-Fidelity DNA Polymerase (New England Biolabs), or repliQa HiFi ToughMix (Quanta Bio). Water was purified using a MilliQ water purification system (MilliporeSigma). Antibiotics (Gold Biotechnology) were used at the following concentrations for plasmid selection, unless otherwise noted: 30 μg/mL kanamycin, 40 μg/mL chloramphenicol, 50 µg/mL carbenicillin, 100 μg/mL spectinomycin, and 10 μg/mL trimethoprim. Antibiotics were used at one-third concentrations for strains bearing three unique plasmids, one-quarter concentrations for strains bearing four unique plasmids, and one-fifth concentrations for strains bearing five unique plasmids. eDRM was prepared by supplementing DRM ^68^ with 6.3 g/L peptone (Millipore), 0.63g/L yeast extract (Thermo Scientific), and 22.5mM oxalate (Combi-Blocks). We note that higher oxalate concentrations can result in significant cell death. Unless otherwise noted, all DNA assembly and plasmid propagation was performed in NEB Turbo cells (New England Biolabs) or Mach1F cells ^37^ (Mach1 T1^R^ cells (Thermo Fisher Scientific)) mated with F’ episome of S2060 ^69^).

### USER cloning

All plasmids were cloned using USER (Uracil-Specific Excision Reagent) assembly. The primers were designed to include a USER junction (spanning from a 5’ terminal dA to a dU base 13-20 base pairs downstream). USER junctions were designed to have a 30°C < T_m_ < 60°C and minimal secondary structures. PCR products were run on a 1% agarose gel in 1x sodium borate buffer ^70^ containing approximately 0.2 µg/mL ethidium bromide, allowing visualization under ultraviolet light. Agarose containing PCR fragments was dissolved in Qiagen QG buffer, purified using DNA purification columns (Bio Basic), and eluted in MilliQ water. Fragments were quantified using a NanoDrop 2000 Spectrophotometer (Thermo Fisher Scientific). For USER assembly 0.1 to 0.5 pmol fragments were added in an equimolar ratio in a 10 μl reaction that included 0.75 µL USER enzyme (Uracil-DNA Glycosylase and DNA-glycosylase-lyase Endonuclease VIII; New England Biolabs), 1ul rCutSmart Buffer (50 mM potassium acetate, 20 mM Tris-acetate, 10 mM magnesium acetate, 100 µg/mL recombinant albumen at pH 7.9; New England Biolabs) and 0.75 µL DpnI (New England Biolabs). The reactions were incubated at 37°C for 20 min, followed by heating to 80°C for 3 min and slow cooling to 12°C ramped at 0.1°C/s in a thermocycler. The USER products were used for heat-shock transformation of chemically competent NEB Turbo or Mach1F cells.

### Chemically competent cell preparation

To prepare competent cells, an overnight culture was diluted 100-fold in 2xYT media supplemented with maintenance antibiotics and grown at 37 °C with shaking at 350 rpm to OD_600_ 0.5–0.7. Cells were pelleted by centrifugation at 5000 × rcf for 5 min at 0°C. The spent media was decanted and the pellet was resuspended in the residual media by keeping it on ice for 10-15 min and gently shaking at short regular intervals. TSS (2xYT medium supplemented with 5% v/v DMSO, 10% w/v PEG 3350, and 20 mM MgCl_2_) at a volume of 10% of the original culture was added to the resuspend cells and mixed by gently swirling. The cell suspension was then aliquoted, flash-frozen in liquid nitrogen, and stored at −80°C until use.

### Transformation of chemically competent cells

To transform cells, 100-400 μL of competent cells were thawed from –80 °C on ice for 15 min. An equal volume of KCM solution (100 mM KCl, 30 mM CaCl_2_, 50 mM MgCl_2_ in H_2_O) was added to the tube and mixed gently (not vortexed). For each transformation, 2 μL of plasmid DNA (up to 3 plasmids per transformation) was added to 25 μL aliquots of competent cells/KCM mix. The mixture was incubated on ice for 10-30 min and heat-shocked at 42°C for 90 seconds. The mixture was then chilled on ice for 2 min, then added to 1 mL of 2xYT media.

Cells were allowed to recover at 37 °C with shaking at 350 RPM for at least 45 min before being streaked on 2xYT agar plates (1.5%) containing the appropriate antibiotics and incubated at 37 °C for 16−18 hours.

### Bacterial genomic modifications

To delete target genes from the *E. coli* genome, a single DNA fragment encoding the KanR–SacB cassette was amplified using primers with 40 base pair homology regions to the gene of interest (HR1 and HR2) ^23^. PCR products were transformed into electrocompetent *E. coli* cells carrying pKD46 ^71^ and pre-induced with arabinose to overexpress the lambda red proteins. The mixture was allowed to recover in 2 ml of 2xYT at 30°C overnight then struck the following morning onto solidified 2xYT + Kanamycin to select for transformants. Isolated colonies were cultured in 2xYT liquid medium with Kanamycin at 30°C overnight. Aliquots of the overnight cultures were diluted 100-fold in water to serve as template for PCR-based genotyping. Primer pairs flanking the deleted region were used to verify mutants with target gene deletion and KanR-specific primers were used to verify KanR insertion. Clones bearing the desired deletions were then made electrocompetent and transformed with oligos encoding only the HR1 and HR2 sequences, recovered at 30°C overnight then struck the following morning onto solidified 2xYT + 5% sucrose to select loss of the KanR–SacB cassette. After 24∼36 hours, single colonies were picked into 100 μl water and an aliquot of the cell suspension was used as a PCR template to verify the deletion with the same pair of flanking primers used to confirm KanR-sacB insertion. A product size transition (typically ∼3kb to ∼400bp) is indicative of the deletion, which was also confirmed by testing for sensitivity to Kan.

### Adaptive Laboratory Evolution (ALE)

To evolve glycolytically disrupted *E. coli* mutants toward optimal growth, adaptive laboratory evolution was carried out to slowly shift the media dependence of the evolving population. A starter culture of S3711 cells was used to seed a chemostat at an initial density (OD_600_) of 0.05 into 250 mL of DRM^A^ supplemented with 40 mM glycerol and 40 mM succinate into duplicate chemostats. The flow rate of fresh DRM^A^ + glycerol/succinate was initially maintained at ≤ 0.75 vol/hr for 7 days to allow the populations to genetically drift. After this period, the DRM titration was initiated at 0.001%, and then doubled to 100% over the course of 22 days. Over the course of the ALE experiment, higher DRM concentrations required reductions of the chemostat flow rate to overcome the poor doubling time of the strain, stabilizing at 0.27 vol/hr at 100% DRM. After reaching 100% DRM, the flow rate was increased from 0.27 vol/hr to 2 vol/hr over 33 days to select for the fittest mutants. Samples from each chemostat were collected daily throughout the evolution campaign and plated on DRM^A^ + glycerol/succinate, DRM complete, and 2xYT to determine the titer of each chemostat and the relative fitness on all media.

### Doubling time analysis and kinetic reporter assays

Typically, four single colonies of each strain or chemostat population were picked into 2xYT, DRM, and M9GS media and grown overnight at 37 °C with shaking at 900 RPM. Saturated cultures were diluted 50-fold into the relevant medium and 200 µl of each culture was transferred to a 96-well black wall, clear bottom plate (Costar) and overlaid with 20ul mineral oil per well. The plate was incubated in a Spark (Tecan) plate reader running SparkControl v2.3 maintained at 37°C with shaking (1440 rpm). Optical density was measured every 10 min over a 12-hour period and the doubling time of each well was calculated individually using GraphPad Prism (version 9).

### CdaR directed evolution

To evolve CdaR toward higher specificity for glycerate, an error-prone PCR library were generated using GeneMorph II Random Mutagenesis Kit (Agilent Technologies) and cloned into a plasmid expressing a tripartite fusion protein that includes barnase, mCherry, and chloramphenicol acetyltransferase (Cat) under control of the CdaR-responsive P_garP_ promoter.

The assembled library was transformed directly into S4012 and plated on 1.5% agar plates supplemented with either 2xYT or DRM alongside the maintenance antibiotics. After ∼36 hours of growth, each plate was scraped to retain all genotypic diversity, diluted to a starting OD_600_ of 0.05, then grown to make chemically compotent cells. Each population was transformed with a barnase overexpression plasmid, recovered for 3 hours at 37 °C with shaking at 350 rpm, then plated on 1.5% agar plates supplemented with either 2xYT or DRM alongside the maintenance antibiotics and 20 µg/mL chloramphenicol. After 48 hours of growth, 48 single colonies were picked from each of the four conditions overnight in 2xYT liquid medium. The following morning, each culture was diluted 50-fold into either either 2xYT or DRM supplemented with the maintenance antibiotics with or without 1 mM glycerate (Millipore Sigma) in a 96-well 2 mL deep well plate, sealed with a porous film, and grown at 37 °C with shaking at 900 rpm for 24 hours. Plates were then quantified as indicated below.

### Reporter assays

S4221, S4009, S4012, or S5643 cells harboring the appropriate biosensing plasmids were transformed alongside plasmids expressing RuBisCO variants and grown on 2xYT (United States Biological) with 1.5% agar supplemented with maintenance antibiotics and incubated at 37 °C overnight. Colonies were picked the following day into liquid super broth (SB) or 2xYT medium (United States Biological; SB shows improved growth over 2xYT) supplemented with the appropriate antibiotics and incubated at 37 °C with shaking at 900 rpm for 18 hours. For fluorescence assays, overnight cultures were diluted 50-fold into liquid DRM or eDRM medium (eDRM outperforms DRM for most assays) supplemented with maintenance antibiotics in a 96-well 2 mL deep well plate, sealed with a porous film, and grown at 37 °C with shaking at 900 rpm for 24 hours. In some cases, 0-100 ng/mL anhydrotetracycline (Honeywell Fluka) was added to the 96-well 2 mL deep well plate if the RuBisCO expression plasmid was under the control of the tetAp promoter. For luminescence assays, overnight cultures were diluted 50-fold into liquid DRM or eDRM medium supplemented with maintenance antibiotics in a 96-well 2 mL deep well plate, sealed with a porous film, and grown at 37 °C with shaking at 900 rpm for 4-6 hours. To quantify output, 150 µL of each culture was aliquoted into a 96-well black wall, clear bottom plate (Costar) after the appropriate incubation. OD_600_ and the excitation and emission wavelengths (fluorescence) or luminescence settings were used for measurements using a Spark (Tecan) plate reader running SparkControl v2.3. Reporter signal was normalized to OD_600_ after blank media subtraction. GraphPad Prism (version 9) was used for plotting and data analysis. Where relevant, non-linear dose-response functions were fitted to each variant’s mScarlet-I signal to determine the EC_50_ of the aTc titration, and Pearson correlation coefficients as well as P values were calculated in Prism. For fluorescence assays, any replicates that did not grow to an OD_600_>1.0 were removed from the subsequent analysis (it is not unusual for one of the four replicates to not grow, or to grow and not fluoresce).

### Diffusion assays

WT and transporter/symporter-deleted S4012 cells were transformed with plasmids encoding Prk, *Tt*HA0368, and either a functional CbbM plasmid + mScarlet-I-based reporter (sender strain) or non-functional CbbM (K191M) plasmid + mEmerald-based reporter (receiver strain). Tranformed cells were plated on 2xYT (United States Biological) with 1.5% agar supplemented with maintenance antibiotics and incubated at 37 °C overnight. Isolated colonies were picked into SB medium supplemented with maintenance antibiotics and grown at 37°C with shaking at 350 rpm overnight. Overnight cultures were diluted 50-fold into eDRM supplemented with maintenance antibiotics in a 96-well 2 mL deep well plate and grown at 37°C with shaking at 900 rpm overnight. For each diffusion assay between plated culture, 1 uL of overnight DRM cultures were spotted on a DRM plate supplemented with maintenance antibiotics and incubated at 37°C. For the glycerate diffusion assay, 1 uL of 50 mM glycerate was spotted adjacent to 1 uL spots of saturated receiver cultures on an eDRM plate supplemented with maintenance antibiotics and incubated at 37°C. Each array of spots contains one spot of the donor strain (or glycerate) at the top left-hand corner and 3 biological replicates of a receiver strain. The fluorescence images were taken using an Azure c400 Imaging system after 48 hours of incubation.

### Mock evolution

An NNK degenerate library randomizing all CbbM essential residues K191, D193 and E194 was created by PCR using primers synthesized to be degenerate at those positions. The PCR was performed using the inactive template CbbM (K191M) to minimize contamination in downstream steps. Next, this fragment library was USER-assembled and transformed into S5643 competent cells harboring *Tt*HA0368, PrkA(R52A), and CdaR^evo1^. Transformants were plated on 2xYT (United States Biological) with 1.5% agar supplemented with maintenance antibiotics and incubated at 37 °C overnight. After 24 hours of growth, all resultant colonies were pooled by scraping, resuspended in eDRM liquid medium, diluted to starting OD_600_ of 0.05, and incubated overnight at 37°C with 5% CO_2_. After 24 hours of growth, the library culture was washed with PBS buffer three times successively then diluted 100-fold in preparation for fluorescence activated cell sorting (FACS). Analysis was carried out using a Beckman Coulter Astrios, with excitation/emission wavelengths of 561/614nm and 488/513nm for mScarlet-I and sfGFP, respectively. The sorting gate was chosen to select for intact cells within the top 4.42% of fluorescence values for mScarlet-I. Collected cells were immediately plated on 2xYT with 1.5% agar supplemented with maintenance antibiotics and incubated at 37 °C overnight. Cultures were used to inoculate an mScarlet-I fluorescence assay as described above. Following confirmation of positive mScarlet-I signal, RuBisCO plasmids from functional cultures were purified and subjected to Sanger sequencing.

### Computational analyses

Structure predictions of *T. thermus* HA0368, *E. coli* CdaR, and *R. rubrum* CA were made using Alphafold ^62,63^ run on Google Collaboratory ^72^. Protein-ligand interactions were predicted using CB-Dock ^64^. The phylogenetic tree was constructed using Clustal Omega ^73^ multiple sequence alignment and iTOL ^74^ for visualization.

## Acknowledgments

The authors would like to thank all members of the Badran Lab for helpful discussions and suggestions. The authors are grateful to Prof. Ron Milo (Weizmann Institute of Science) for kindly sharing their library of RuBisCO variants, and as well as Prk and CA expression plasmids.

## Author Contributions

DLL designed the study, performed experiments, analyzed results, and wrote the paper. ZL designed the study, performed experiments, and analyzed results. AHB conceived of the concept, designed the study, performed experiments, analyzed results, supervised the research, and wrote the paper.

## Data and materials availability

All data are available in the main text or the extended data, and raw data generated in this manuscript are available upon request. Correspondence and requests for materials should be addressed to. Key materials required to use this system have been deposited in Addgene

